# Contesting the evidence for gene expression in lower oceanic crust

**DOI:** 10.1101/2020.03.25.005033

**Authors:** William D. Orsi

## Abstract

Li *et al* [*Nature* **579**, 250–255 (2020)] report microbial gene expression in deep ocean crust, from microbial communities that have concentrations as low as 100 cell per cubic cm. This would be incredible, since that would be the lowest biomass sample ever analyzed for gene expression for a deep biosphere sample by several orders of magnitude. However, I have reanalyzed the data and show that the author’s data are derived from contamination via seawater, humans, human skin associated bacteria, drilling fluid, and molecular reagent kits. The method that the authors used to produce their gene expression data is highly sensitive to DNA contamination, which is the case here. The authors claim that *Pseudomonas*, a known widespread contaminant in molecular reagent kits, is one of the most active groups in their lower crust samples and base their metabolic analysis on this contaminating organism. Here, I show that the gene expression data are derived from contaminating *Pseudomonas* (as well as other groups), and I show that the groups claimed as being active in the rock samples are also present in the negative controls at similar or higher relative abundance. For example, methane production in long term incubations was reported but gene expression from methanogens was 10 times higher in the drill fluid negative controls compared to the rock samples, demonstrating that the methane measured is likely derived from methanogen contaminants introduced from the drill fluid. There are no signs of active life here, but lipid biomarkers were found preserved in the rocks. Thus, the authors have sampled a fossilized biosphere, not a living one.

## Main Text

Obtaining authentic measurements of microbial activity in subsurface habitats is challenging because the cell concentrations and the metabolic rates per cell are extremely low, and the risk of contamination is very high. Li *et al*.^1^ reported gene expression from a microbial biosphere in the lower ocean crust, from samples that have cell concentrations as low as 100 cells per cm^−3^. This would be incredible, since that would be the lowest biomass sample ever analyzed for gene expression for a subsurface sample by several orders of magnitude.

The authors discuss *Pseudomonas* as being one of the most active groups in their samples^1^, which is concerning because *Pseudomonas* is one of the most common contaminants in molecular reagent kits^2^. This prompted me to check the annotations of the entire dataset provided by the STAT annotation pipeline using the NCBI RefSeq database available on the short read archive (SRA) website. This shows that sequences annotated as human are abundant in all metatranscriptomes of Li *et al*, up to 70% (Figure 1A). The remaining sequences are annotated as Bacteria typically found on human skin, and those commonly found in kit contaminants (e.g., *Acinetobacter, Streptococcus, Staphylococcus*) (Figure 1A). This suggests that the metatranscriptomes are heavily contaminated. However, the RefSeq database used for the default annotation pipeline by the SRA only includes high quality completed genomes, and does not include the numerous metagenome assembled genomes (MAGs) and single cell genomes (SAGs) from subsurface samples.

**Figure 1:**
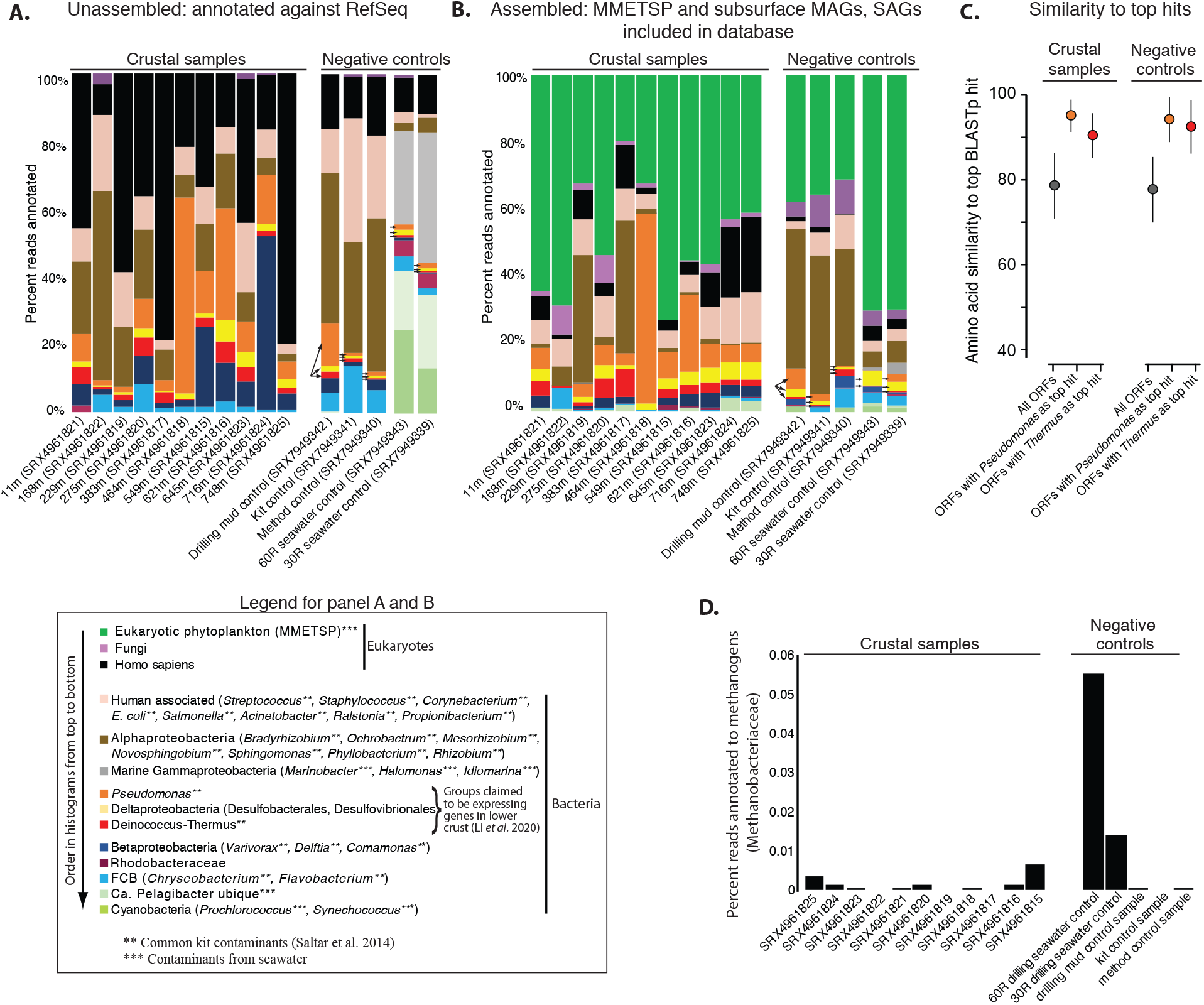
Comparison of lower crust metatranscriptomes to negative controls. The data are summarized showing the relative abundance of taxa that were the top BLAST hit for unassembled (raw reads) compared against the RefSeq database (A), and assembled data compared against the RefSeq database with the MMETSP and subsurface MAGs and SAGs added (B). The legend shows the genera that were included into each category, note the abundance of seawater and human contamination in the samples. Arrows show the sequences with hits to groups in the negative controls, that were claimed by Li *et al* as expressing genes in the lower crustal samples. Panel C shows high similarity of the *Pseudomonas* and *Thermus* open reading frames to their top hit in the RefSeq database, which was not different between samples and controls. Panel D shows that the two of the seawater drilling fluid negative controls actually have the highest relative abundance of transcripts methanogen groups.

Thus, I double checked the annotations of the authors data against a larger database^3^ that in addition to RefSeq includes subsurface MAGs and SAGs including all those from ocean crustal fluids^4,5^, as well as all of the transcriptomic data from microbial eukaryotes produced in the Marine Microbial Eukaryote Sequencing Project^6^. This did not result in an increased annotation of sequences from typical subsurface groups, but instead revealed that most of the authors data are actually derived from photosynthetic eukaryotic algae from seawater (e.g., diatoms, chlorophytes, and dinoflagellates), where they make up >50% of the data in all the authors samples (Figure 1B). The remaining sequences have highest similarity to Humans, and Bacteria typically found on human skin and kit contaminants like *Pseudomonas, Streptococcus, and Staphylococcus.* The domination of sequences from photosynthetic eukaryotes that are plankton that live in the surface ocean, indicate that the metatranscriptomes of Li *et al* are heavily contaminated with seawater. Indeed, the 16S rRNA gene data from Li *et al* are dominated by bacterial groups common to seawater. A bacterial cell in the sunlit and relatively nutrient rich seawater has on average 200-2000 mRNA molecules per cell^7^, and levels in subsurface microbes should be even lower given the extreme energy limitation^8^. In contrast, eukaryotic phytoplankton have relatively large genomes and a single cell can have up to 50,000 mRNA molecules^9^, and thus it would only take nucleic acids from small number of contaminating eukaryotic phytoplankton from seawater to swamp an entire metatranscriptome sequencing run on such low biomass samples.

The high amount of seawater contamination is exacerbated by the mRNA amplification method used by Li *et al*^1^. The Ovation RNA-Seq kit that the authors used has a high fidelity SPIA polymerase for amplifying nucleic acids, and is highly sensitive to DNA contamination when used on minute amounts of RNA^10^. Such low biomass samples would be best worked with in an ISO Class 4 clean room as common for single cell genomics labs. But, the authors instead worked in HEPA filtered laminar flow hoods, which help to reduce contamination but still can be easily contaminated by a few single contaminating human skin cells and or aerosols containing nucleic acids. In an attempt to increase the amount of cells, the authors used 40g of rock material – but this does not solve the problem since the amount of extraction buffer needed to process the samples needs to be scaled up accordingly which in turn concentrates any contaminants in the extraction buffer. Considering this, it is understandable that processing such low biomass samples in this manner would be easily lead to contamination from seawater, aerosols, human skin, or drilling fluid. Indeed, many of the the groups claimed as active by Li *et al* in the samples are also presented in all negative control samples (Figure 1A, B).

Li *et al*^1^ write that they took extensive measures to reduce contamination during sampling, and performed negative controls in the form of extraction blanks of DNA and RNA. They write in the Methods that they removed sequences from “…taxa known to be common molecular kit reagent contaminants^46,56^”, which in the cited Salter *et al*^2^ paper includes Deinococcus-Thermus and *Pseudomonas*. Curiously, the authors then proceed to discuss sequences from Deinococcus-Thermus and *Pseudomonas* as endemic to their samples even though they admit in the Methods that these are common kit contaminants. Considering this, it is not surprising that Deinococcus-Thermus, *Pseudomonas,* and sulfate reducing Deltaproteobacteria were represented in all of the negative controls (Figure 1A, B) which indicates they were introduced as contaminants. The high amount of seawater contamination further demonstrates this is a strong possibility The authors allude to the possibility that their sequences are new groups of Deinococcus-Thermus and *Pseudomonas* not yet in databases, but the average similarity of these sequences are >90% to their top hits, which is even higher compared to the median value (Figure 1C). This shows that the sequences from Deinococcus-Thermus and *Pseudomonas* in the authors metatranscriptomes are not from novel species but rather contaminating organisms from the drill fluid, seawater, and the kits themselves (Figure 1A, B). The metabolic pathway analysis provided by the authors focuses on conserved metabolism (e.g., amino acid biosynthesis and degradation, Krebs cycle) that are widespread in contaminating bacteria from reagent kits^2^, including Deinococcus-Thermus and *Pseudomonas*.

The authors then took these highly contaminated datasets, and attempt to remove contaminants by co-assembling them with the sequences deriving from the five control samples, claiming that this “…affords the highest possible confidence about any transcripts discussed.” But, I show here that this type of approach easily misses contaminant sequences. For example, transcripts from both the lower crust samples and the negative controls map to the 16S and 23S rRNA genes of *Pseudomonas fluorescens* with high coverage, indicating this organism is a contaminant (Figure 2). However, just downstream of the ribosomal operon in the same contaminant genome the transcripts from the lower crust samples and negative controls map to two separate genes and do not co-assemble (Figure 2). This does not afford high confidence about the transcripts as the authors claim, but does explain how such contaminating genes remained in the authors dataset after the quality control measures used by Li *et al*.

**Figure 2:**
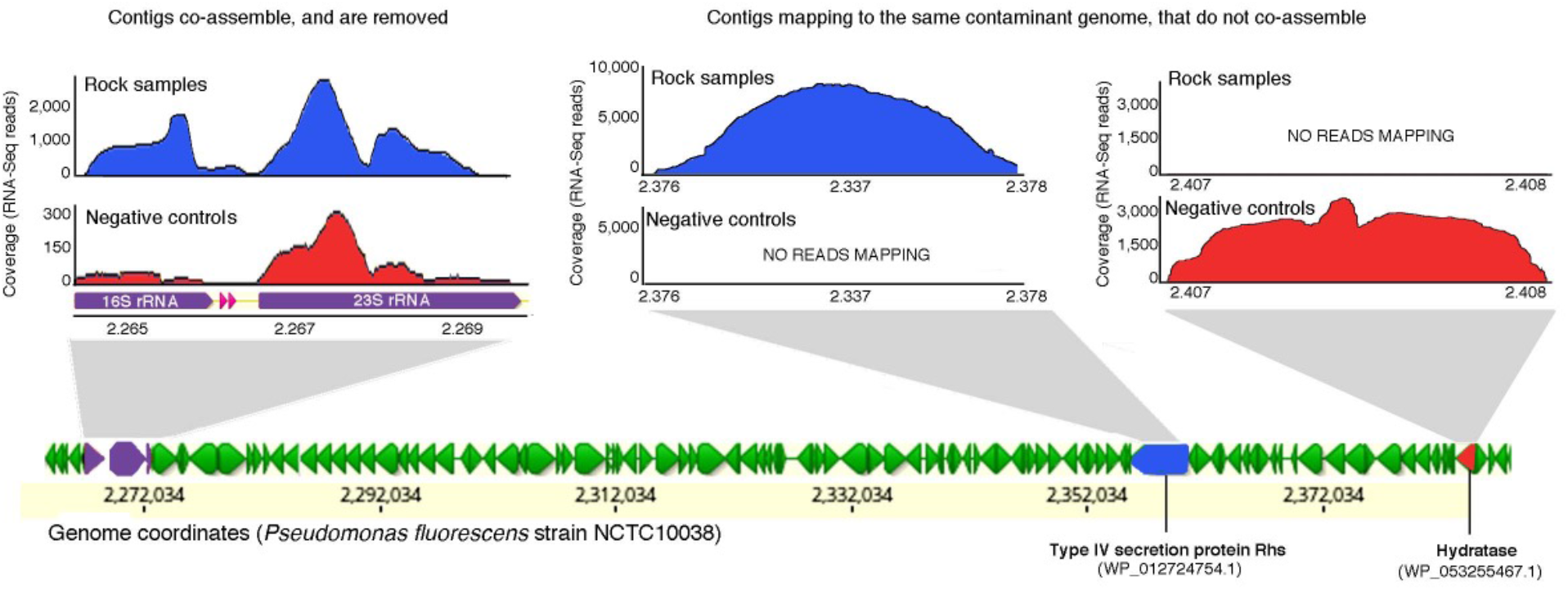
Contigs from samples and controls that map to the same contaminant genome do not co-assemble. Transcripts from the negative controls and the samples both map to the 16S and 23S rRNA subunits of *Pseudomonas fluorescens* with high stringency (>95% similarity) indicating this is a contaminant. However, reads mapping to the type IV secretion protein Rhs and hydratase do not co-assemble despite being derived from the same contaminant genome. The contaminant removal method used by Li *et al* would miss such contaminant sequences.

But what about the methane measured by Li *et al* in the long term incubations? The authors argue that methane formation in incubations is indication of an *in situ* active community. But, the metatranscriptomes from drill fluid negative controls have gene expression from methanogens (family Methanobacteriaceae), that was 2-10 times higher compared to the rock samples (Figure 1D). This strongly indicates that the methane production in the incubation likely stems from contaminating methanogens in the Methanobacteriaceae that were introduced from the drilling fluid. This is understandable, considering that methanogens in the genus *Methanobrevibacter* (Methanobacteriaceae) are a typical contaminant in scientific drilling cores that can derive from anthropogenic wastewater ^11^. Li *et al* obtained their samples via rotary coring, and the drill fluid used that contained the active methanogens (Figure 1D) could have easily been introduced into or onto the samples during drilling.

Despite being included in the database (Figure 1B), sequences annotated to MAGs and SAGs from inhabitants of crustal subsurface habitats such as ammonia oxidixing Thaumarchaeota, Bathyarchaeota, sulfide oxidizing Epsilonproteobacteria, and iron oxidizing Zetaproteobacteria^4,5,12^ are notably missing in the gene expression dataset of Li *et al*^1^ (Figure 1B). While all subsurface gene expression data are prone to contamination due to the low biomass as an unavoidable challenge, it is nevertheless expected in the case of truly active subsurface microbial communities that at least some sequences derived the typical inhabitants of these extreme environments should be represented with reliable levels of sequencing coverage. Sequences from sulfate reducing Deltaproteobacteria are proposed by Li *et al*^1^ as also being endemic, which is plausible since they were annotated in both analyses (Figure 1), are strict anaerobes, and are not typical contaminants from kits or human skin. But, the same sulfate reducing Deltaproteobacteria groups are also present in the negative controls including the drill fluid and seawater (Figure 1A, B). This does not support the interpretation that these sequences derive from living sulfate reducing bacteria expressing genes in the lower crust, but are instead derived from contaminants.

The high amount of contamination from seawater, human skin, and reagent kits, as well as the presence of transcripts from *Pseudomonas*, *Thermus*, methanogens, and Deltaproteobacteria in the drill fluid controls (Figure 1) calls the conclusion of a living biosphere in the lower crust into question. Moreover, because co-assembly methods of removing contaminant sequences misses contaminant sequences (Figure 2), the transcripts passing quality control by Li *et al* cannot be trusted as endemic. While there are no signs of active life in these lower crust samples, lipid biomarkers were found preserved in the rocks as well as photographic evidence of preserved biomass structures that resemble those seen previously in ocean crust^13^. Thus, it appears that the authors have sampled a fossilized biosphere, not a living one.

## Methods

Annotation summaries of the raw reads were downloaded from the SRA website, available from the NCBI SRA Taxonomy Analysis Tool ^14^ which uses the RefSeq database and is the default annotation pipeline available on the SRA (https://github.com/ncbi/ngs-tools.git). Quality controls, *de novo* assembly, and ORF finding were performed as described previously^3^. Reads were mapped against the *P. fluorescens* genome using Geneious 20.0.4. ORF annotations were performed as described previously using a database that in addition to RefSeq includes subsurface MAGs and SAGs including all those from ocean crustal fluids^3^, as well as all of the transcriptomic data from microbial eukaryotes produced in the Marine Microbial Eukaryote Transcriptome Sequencing Project (MMETSP)^6^. The predicted proteins from the MMETSP used in this analysis can be downloaded here (https://github.com/wrf/misc-analyses/tree/master/marine_meta).

